# Benefits of higher resistance-training volume depends on ribosome biogenesis

**DOI:** 10.1101/666347

**Authors:** Daniel Hammarström, Sjur Øfsteng, Lise Koll, Marita Hanestadhaugen, Ivana Hollan, William Apro, Jon Elling Whist, Eva Blomstrand, Bent R. Rønnestad, Stian Ellefsen

## Abstract

Resistance-exercise volume is a determinant of training outcomes. However not all individuals respond in a dose-dependent fashion. In this study, 34 healthy individuals (males *n =* 16, age 23.6 (4.1) years; females *n =* 18, 22.0 (1.3) years) performed moderate- (3 sets per exercise, MOD) and low-volume (1 set, LOW) resistance training contralateral fashion for 12 weeks (2-3 sessions × week^−1^) enabling intra-individual comparisons of effects of training modalities. Muscle cross-sectional area (CSA) and muscle strength was assessed at weeks 0 and 12, along with biopsy sampling (m. Vastus lateralis). Muscle biopsies were also sampled before and one hour after the fifth session (Week 2). MOD resulted in larger increases in muscle CSA (5.2 (3.8)% versus 3.7 (3.7)%, *P* < 0.001) and strength (3.4-7.7% difference, all *P <* 0.05). In muscle, this coincided with greater reductions in type IIX fibres from week 0 to week 12 (MOD, −4.6 vs. LOW - 3.2%-point), greater post-exercise (Week 2) phosphorylation of mTOR (12%), S6-kinase 1 (19%) and ribosomal protein S6 (28%, Week 2), greater rested-state total RNA (8.8%, Week 2) and greater exercise-induced elevation of c-Myc mRNA expression (25%, Week 2; all *P <* 0.05). Fifteen participants displayed robust benefits of MOD on muscle hypertrophy. This was associated with greater accumulation of total RNA at Week 2 in MOD vs. LOW as every 1% difference increased the odds of MOD benefit by 5.4% (*P* = 0.010). In conclusion, MOD led to on average greater adaptations to resistance training and dose-dependent hypertrophy was associated with volume-dependent regulation of total RNA at week 2. This suggests that ribosomal biogenesis regulates the dose-response relationship between training volume and muscle hypertrophy.

**Key points:**

- For individuals showing suboptimal adaptations to resistance training, manipulation of training volume is a potential measure to facilitate responses. This remains unexplored in previous research.
- Here, 34 untrained individuals performed contralateral resistance training with moderate and low volume for 12 weeks. Overall, moderate volume led to larger increases in muscle cross-sectional area, strength and type II fibre-type transitions.
- These changes coincided with greater activation of signaling pathways controlling muscle growth and greater induction of ribosome synthesis.
- Fifteen individuals displayed clear benefit of moderate-volume training on muscle hypertrophy. This coincided with greater total RNA accumulation in the early-phase of the training period, suggesting that ribosomal biogenesis regulates the dose-response relationship between training volume and muscle hypertrophy.
- These results demonstrate that there is a dose-dependent relationship between training volume and muscle hypertrophy. On the individual level, benefits of higher training volume was associated with increased ribosomal biogenesis.

## Introduction

In humans, the biological adaptation to resistance training varies with exercise-training variables such as volume, intensity, rest between repetitions and sets, selection and order of exercises, repetition velocity and frequency of training sessions (Ratamess *et al*., 2009), as well as with genetic and epigenetic disposition and environmental factors (Timmons, 2011; Seaborne *et al*., 2018; Morton *et al*., 2018). As time constraints often hinder participation in exercise training-programs (Choi *et al*., 2017), numerous studies have searched for the minimally required exercise dose to promote beneficial adaptations. Within-session volume has received particular attention, and indeed, a handful studies have shown that low-volume training provides similar gains in strength and muscular mass as moderate-volume training (Cannon & Marino, 2010; Ostrowski *et al*., 1997; Mitchell *et al*., 2012), though meta-analyses conclude in favor of moderate volume protocols (Rhea *et al*., 2003; Krieger, 2009, 2010; Schoenfeld *et al*., 2016). This apparent failure of specific studies to disclose benefits of increased training volume is likely due to a combination of small sample sizes and substantial variation in training responses between individuals and experimental groups. In theory, within-participant designs should alleviate these limitations.

Individual response patterns to resistance training, including muscle strength and mass, correlate closely with muscle cell characteristics, measured in both rested-state and acute training-phase conditions (Thalacker-Mercer *et al*., 2013; Stec *et al*., 2016; Terzis *et al*., 2008). Of particular interest is the molecular signatures conveyed by the mechanistic target of rapamycin complex 1 (mTORC1) and its associated downstream target S6 kinase 1 (S6K1). This pathway acts as a master signaling hub of muscle fiber hypertrophy by controlling protein synthesis and degradation (Laplante & Sabatini, 2012). Inhibition of mTORC1 signaling impairs protein synthesis in humans (Drummond *et al*., 2009), and exercise-induced activation of mTORC1 signaling correlate with increase in muscle protein synthesis and subsequent muscle growth (Burd *et al*., 2010; Terzis *et al*., 2008). In line with this, surplus training volume leads to greater phosphorylation of S6K1 (Burd *et al*., 2010; Terzis *et al*., 2010; Ahtiainen *et al*., 2015), and increased myofibrillar protein synthesis (Burd *et al*., 2010), fitting the notion that increased training volume provides more pronounced adaptations. However, also from a cell biological perspective, present findings on effects of different training volumes are heterogeneous. For example, Mitchell *et al*. (2012) failed to show differences in S6K1 phosphorylation between volume protocols, corroborating with similar effects of different volumes on muscle strength and mass.

In muscle cells, increased mTORC1 activity leads to increased translational efficiency through activation of 4E-BP1 and S6K1 (Laplante & Sabatini, 2012). It also leads to increased translational capacity, measured as de novo ribosomal biogenesis controlled synergistically with mTORC1 by c-Myc activity and subsequent transcription of ribosomal RNA (rRNA) (Nader *et al*., 2005; West *et al*., 2016). Recent observations in humans indicate that translational capacity is a limiting factor for training-induced muscle hypertrophy. First, increased abundances of rRNA in response to resistance training, measured as total RNA per-weight-unit muscle tissue, correlate with muscle hypertrophy (Figueiredo *et al*., 2015). In accordance with this, training-induced increases in rRNA are larger in high-responders than in low-responders (Stec *et al*., 2016; Mobley *et al*., 2018). Second, elderly typically show blunted ribosome biogenesis, coinciding with attenuated hypertrophic responses (Stec *et al*., 2015; Brook *et al*., 2016). Collectively, these observations suggest that muscle growth depends at least in part on increased translational capacity, making it a prime candidate for explaining the diverse response patterns seen to resistance training with different volume in different individuals. To date, no study has investigated the association between training volume, ribosome biogenesis and regulation, and gross training adaptations.

Muscle fibre composition is another potential determinant of muscular responses to resistance training. Type II fibres have greater growth potential compared to type I fibres (Stec *et al*., 2016; Jespersen *et al*., 2011), and readily switch from IIX to IIA phenotypes in response to mechanical loading (Andersen & Gruschy-Knudsen, 2018; Widrick *et al*., 2002; Ellefsen *et al*., 2014*b*), suggesting that these fibers display greater plasticity in response to resistance training.

The purpose of the present study was to evaluate the effects of single- and multiple-sets training protocols on strength, muscle hypertrophy and fibre-type composition using a within-participant design. In addition, phosphorylation of proteins in the mTORC1 pathway as well as total and ribosomal RNA were determined.

## Methods

### Ethics statement

All participants were informed about the potential risks and discomforts associated with the study and gave their informed consent prior to study enrollment. The study design was pre-registered (ClinicalTrials.gov Identifier: NCT02179307), approved by the local ethics committee at Lillehammer University College, Department of Sport Science (nr 2013-11-22:2) and all procedures were performed in accordance with the Declaration of Helsinki.

### Participants and study overview

Forty-one male and female participants were recruited to the present study with eligibility criteria’s being non-smoking and age between 18 and 40 years. Exclusion criteria were intolerance to local anesthetic, training history of more than one weekly resistance-exercise session during the last 12 months leading up to the intervention, impaired muscle strength due to previous or current injury, and intake of prescribed medication that could affect adaptations to training. During data analyses, seven participants were excluded due to not completing at least 85% of the scheduled training sessions with reasons being: discomfort or pain in the lower extremities during exercise (n=5), injury not related to the study (n=1), failure to adhere to the study protocol (n=1). At baseline, there were no differences in maximal voluntary contraction (MVC) normalized to lean body mass or anthropometrics between included and excluded participants (see Table 1). Among the included group, one participant choose to refrain from biopsy and blood sampling at week 2. Additionally, blood was not collected from three of the participants at different time-points due to sampling difficulties.

**Table 1:** Participant characteristics.

|  | Female |  | Male |  |
| --- | --- | --- | --- | --- |
|  | Included | Excluded | Included | Excluded |
| N | 18 | 4 | 16 | 3 |
| Age (y) | 22.0 (1.3) | 22.9 (1.6) | 23.6 (4.1) | 24.3 (1.5) |
| Weight (kg) | 64.4 (10.4) | 64.6 (9.7) | 75.8 (10.7) | 88.2 (22.4) |
| Stature (cm) | 168 (7) | 166 (8) | 183 (6) | 189 (5) |
| Body fat (%) | 34.1 (5.6) | 28.8 (8.7) | 20.4 (6.0) | 24.3 (15.3) |
| MVC ($\text{Nm} \times \text{LBM}^{-1}$) | 4.9 (0.7) | 5.3 (0.4) | 4.7 (0.8) | 5.0 (0.2) |
| Data are means and (SD) |  |  |  |  |

The intervention consisted of 12 weeks of full-body resistance training (all participants commenced the trial during September-November). Leg-exercises were performed unilaterally to allow within-participant differentiation of training volume. Accordingly, for each participant, the two legs were randomly assigned to perform resistance exercises consisting of one set (single-sets condition) and three sets (multiple-sets condition); i.e. each participant performed both protocols. Muscle strength was assessed at baseline, during and after the training intervention. Body composition was measured before and after the training intervention. Muscle biopsies were sampled from both legs (vastus lateralis) at four time points during the intervention: at baseline (Week 0, rested state), before and one hour after the fifth training session (Week 2 Pre-exercise, rested; Week 2 Post-exercise, acute-phase biopsy) and after completion of the intervention (Week 12, rested state). For overview of the study protocol, see Figure 1.

**Figure 1:**
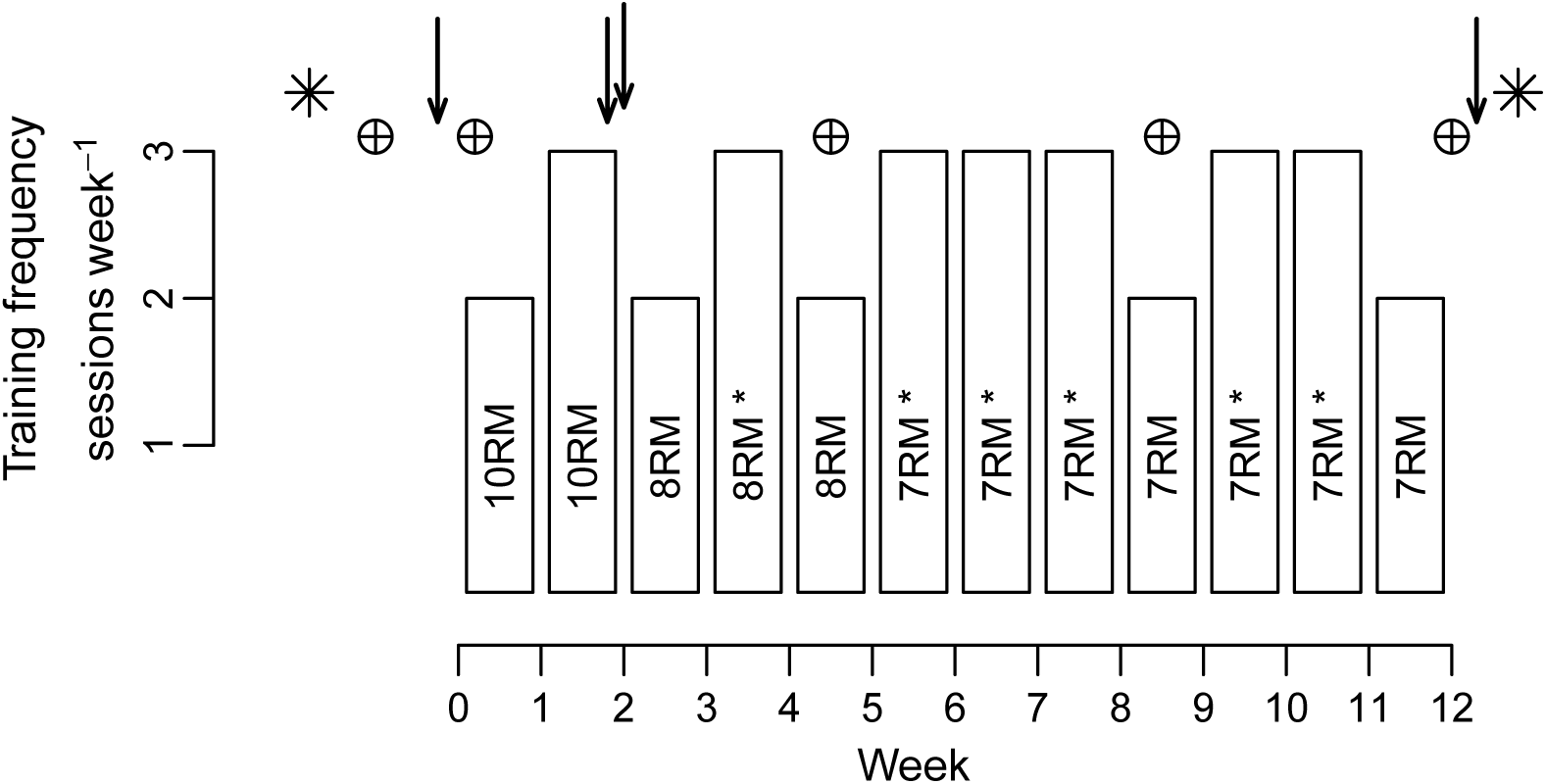
Study overview. Bars represent weekly training frequency with training intensity expressed as repetition maximum (RM). * indicates that one session per week was performed at 90% of prescribed RM intensities. ↓ indicates muscle biopsy: Before (Week 0, n=34) and after the 12-wk intervention (Week 12, n=34), as well as before and after (1h) the fifth exercise session (Week 2 Pre-Ex and Post-Ex, n=33). ⊕ indicates strength test: before the intervention (Week 0, n=34), after 5 and 9 weeks of training (n=18), and after finalization of the intervention (Week 12, n=34). Baseline strength was determined as the highest value obtained during two test sessions performed prior to the intervention. Body composition was measured prior to the intervention (Week 0) and after its finalization (Week 12, n=34) using full-body DXA and knee-extensor muscle MRI (*).

### Resistance-exercise training protocol

Prior to all training-sessions, participants performed a standardized warm-up routine consisting of i) 5-min ergometer cycling (RPE 12-14), followed by ten repetitions each of bodyweight exercises (push-ups with individually adjusted leverage, sit-ups, back-extensions and squats), and iii) one set of ten repetitions at ~50% of 1RM for each of the resistance exercise. Leg resistance exercises were performed in the following order: unilateral leg-press, leg-curl and knee-extension, performed as either one set (single-sets) or three sets (multiple-sets) per exercise. Single-sets were performed between the second and third set of the multiple-sets protocol. Following leg-exercises, participants performed two sets of bilateral bench-press, pull-down, and either shoulder-press or seated rowing (performed in alternating sessions). Rest periods between sets were 90-180 seconds. Training intensity was gradually increased throughout the intervention, starting with 10 repetitions maximum (10RM) the first two weeks, followed by 8RM for three weeks and 7RM for seven weeks (Figure 1). To better fit the training program to participants daily schedule, some sessions were performed unsupervised. The average number of supervised sessions were 91% (SD = 10%, range: 67-100%) of performed sessions. From the ninth training session, every week (containing three training sessions) had one session with reduced loads, corresponding to 90% of the previous session with the same target number of repetitions. Training sessions with maximal effort were separated by at least 48 h. Training sessions with submaximal efforts (90%) were separated from other sessions by at least 24 h. To aid immediate recovery, a standardised drink were given after each session containing 0.15 *g* × *kg*^−1^ protein, 11.2 *g* × *kg*^−1^ carbohydrates and 0.5 *g* × *kg*^−1^ fat.

### Muscle strength assessments

Isokinetic and isometric unilateral knee-extension strength was assessed in a dynamometer (Cybex 6000, Cybex International, Medway USA). Participants were seated and secured in the dynamometer with the knee joint aligned with the rotation axis of the dynamometer. Maximal isokinetic torque was assessed at three angular speeds (60°, 120° and 240° × sec^−1^). Prior to testing, participants were familiarized with the test protocol by performing three submaximal efforts at each angular speed. Participants were given two attempts at 60° × sec^−1^ and three attempts at 120 and 240° × sec^−1^ performed in immediate succession. The highest value was used for statistical analyses. After isokinetic testing, maximal voluntary contraction torque (MVC) was assessed at a knee angle of 30° (full extension = 90°). Participants were instructed to push with maximal force against the lever for 5 sec. Participants were given two attempts, with 30 sec rest in-between. The highest value was used for downstream analyses.

Maximal strength was assessed as one repetition-maximum (1RM) in leg-press and knee-extension. The test session for each exercise started with specific warm-up consisting of ten, six and three repetitions at 50, 75 and 85% of the anticipated maximum. Thereafter, 1RM was found by increasing the resistance progressively until the weight could not be lifted through the full range of motion. For each exercise, the highest load successfully attempted was defined as 1RM. Each participant was given four to six attempts. Prior to the intervention, 1RM was tested twice separated by at least four days with the maximum from the two sessions recorded as baseline 1RM. A subset of the participants (n=18) performed strength assessment during the course of the study (at week 5 and 9). For the remaining participants, ordinary training sessions were prioritized when participants missed out on training or testing due to e.g. illness or scheduling difficulties.

### Muscle cross-sectional area (CSA) and body composition

Knee-extensor muscle CSA (vastus lateralis, medialis, intermedius and rectus femoris) was determined before and after the training intervention using magnetic resonance imaging (MRI) in accordance with manufacturer’s protocol (S-Scan, Esaote Europe B.V., Maastricht, Netherlands). Images were analyzed in a blinded fashion by the same investigator, using OsiriX (v.5.6, Pixmeo Sarl, Bernex, Switzerland). For each participant, CSA was determined at the same distance from the knee-joint pre- and post-intervention (mid-thigh), using at least four consecutive images (5 mm thickness, 10 mm separation; see Figure 2A for representative images). Body composition was determined before and after the intervention using dual-energy X-ray absorptiometry (DXA) (Lunar Prodigy, GE healthcare), in accordance with standard protocol. Prior to MRI and DXA measurements, participants were asked to stay fasted for 2 h and to refrain from vigorous physical activity for 48 h.

**Figure 2:**
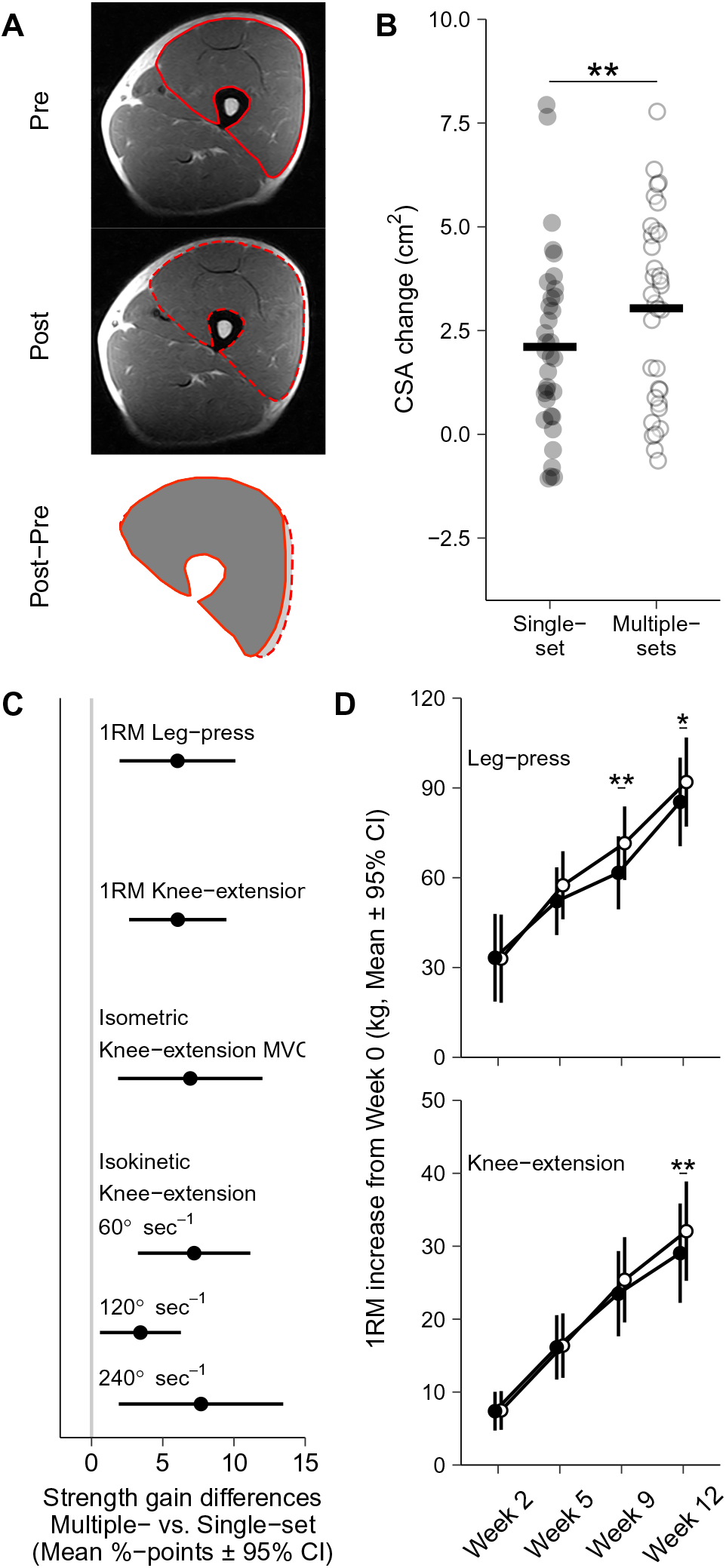
Volume-dependent effects on muscle mass and strength. Training volume-dependent changes in muscle mass and strength after 12 weeks of resistance training, evident as larger increases in knee-extensor muscle CSA measured using MRI (A and B) and larger increases in one-repetition maximum (1RM) knee-extension and leg-press and isometric isokinetic knee-extension strength (C). Time course of changes in 1RM strength (n=18), showing that the difference between volumes occurred towards the end of the training intervention (D). Values are means in B, mean ± 95% CI in C and mean ± 95% CI in D. * represents significant effect of volume-condition * - * * for P<0.05 - P<0.01.

### Hormonal measurements

Hormone analyses were performed on blood samples collected at five time points: along-side muscle biopsies (Figure 1, four sampling events) and 10 minutes after completion of the fifth training session. Samples were drawn from an antecubital vein into serum-separating tubes and kept at room temperature for 30 min before centrifugation (1500 g, 10 min). Serum was immediately aliquoted and stored at −80°C until further processing. Serum concentrations of total testosterone, cortisol, growth hormone and insulin-like growth-factor 1 (IGF-1) were measured on an Immulite 1000 analyzer, using kits from the Immulite Immunoassay System menu (Siemens Medical Solutions Diagnostics, NY, USA), performed according to manufacturer’s protocols. Serum Vitamin D (S-25-OH-D) levels were measured in samples collected before and after the intervention using a electrochemiluminescence immunoassay (Roche Cobas Vitamin D total assay, Roche Diagnostics GmbH., Mannheim, Germany) using automated instrumentation (Roche Cobas 6000’s module e601, Roche Diagnostics GmbH., Mannheim, Germany).

### Muscle tissue sampling and processing

Muscle biopsies were obtained bilaterally from m. vastus lateralis under local anesthesia (Xylocaine, 10 *mg* × *ml*^−1^ with adrenaline 5 *µg* × *ml*^−1^, AstraZeneca AS, Oslo, Norge) using a 12-gauge needle (Universal-plus, Medax, San Possidonio, Italy) operated with a spring loaded biopsy instrument (Bard Magnum, Bard, Rud, Norway). For each participant, resting samples were collected at the same time of day at all time-points and all sampling was done in the morning after a standardised breakfast. Participants were instructed to standardise meals during the last 24 h leading up to the sampling and to refrain from strenous physical activity the last 48 h.

Samples were obtained within 10 minutes from both legs at all time-points. The first biopsy was sampled 1/3 of the distance from the patella to anterior superior iliac spine, subsequent biopsies were sampled ~2 cm proximal from the previous sample. The tissue was quickly dissected free of blood and visible connective tissue in ice-cold sterile saline solution (0.9% NaCl). Samples for immunohistochemistry (~15 *mg*) were transferred to a 4% formalin solution for fixation 24-72 h, before further preparation. Samples for protein and RNA analyses (~60 *mg*) were blotted dry, snap-frozen using −80°C isopentane and stored at −80°C until further analyses.

### Immunohistochemistry

Formalin-fixed muscle biopsies were processed for 2.5 h using a Shandon Excelsior ES (Thermo Scientific, USA), paraffin-embedded and sectioned into 4 *µm* transverse sections. For determination of muscle fibre types, sections were double-stained using BF-35 (5 *µg* × *ml*^−1^, Developmental Studies Hybridoma Bank, deposited by Schiaffino, S.) and MyHCSlow (1:4000, catalog M8421L, Sigma-Aldrich Norway AS, Oslo, Norway). The primary staining was visualized using BMU UltraView DAB and UltraView Red (Ventana Medical Systems, Inc. Tucson, USA). Muscle fibres were counted as either Type I (red), Type IIA (brown), Type IIX (unstained) or hybrid fibers Type IIA/IIX (light-brown) (for representative image, see Figure 5B). Fibres identified as hybrid fibers were analyzed as 0.5 × Type IIA and 0.5 × Type IIX.

### Protein extraction and immunoblotting

Aliquots of muscle-tissue (approximately 25 mg wet weight) were homogenised using a plastic pestle in ice-cold lysis buffer (2 mM HEPES pH 7.4, 1 mM EDTA, 5 mM EGTA, 10 mM MgCl_2_, 1% Triton X-100) spiked with protease and phosphatase inhibitors (Halt, Thermo Fischer Scientific, Life Technologies AS, Oslo Norway), incubated at 4° for 1 hr and centrifuged for 10 min at 10 000 g and 4°C, after which the supernatants were collected. Total protein concentrations were determined on a 1:10 dilution (Pierce Detergent Compatible Bradford Assay Reagent, Thermo Fischer Scientific). The remaining supernatant was diluted to 1.5 *µg* × *µl*^−1^^ total protein in lysis buffer and 4X Laemmli sample buffer (Bio-Rad Laboratories AB, Oslo Norway) containing 2-Mercaptoethanol. Samples were heated to 95°C for 5 min and stored at −20°C until further processing. During analyses, protein samples (20 *µg* of total protein) were separated at 300 V for 30 min using 4-20% gels (Criterion TGX, Bio-Rad), followed by wet transfer to PVDF membranes (0.2 *µm* Immun-Blot, Bio-Rad) at 300 mA for 3 h. Gel electrophoresis and protein transfer were performed at 4°C. Membranes were then stained using a reversible total protein stain (Pierce Reversible Protein Stain, ThermoFischer Scientific) to ensure appropriate protein transfer. Membranes were blocked for 2 h in tris-buffered saline (TBS, 20 mM Tris, 150 mM NaCl) containing 3% bovine serum albumin and 0.1% Tween-20, followed by over-night incubation with primary antibodies targeting either the phosphorylated or non-phosphorylated epitope diluted in blocking buffer followed by 2 h incubation with secondary, horseradish peroxidase-conjugated antibodies diluted in TBS containing 0.1% Tween-20 and 5% skimmed milk. Membranes were washed in TBS containing 0.1% Tween-20 for 6 × 5 min after incubation with primary antibody, and for 8 × 5 min after incubation with secondary antibodies. After chemiluminescent detection (SuperSignal™ West Femto Maximum Sensitivity Substrate, ThermoFischer Scientific), membranes were incubated with hydrogen peroxide (15 min, 37°C) to inactivate the horseradish peroxidase (HRP), as described by Sennepin *et al*. (2009), followed by over-night incubation with primary and secondary antibodies as described above. If the phosphorylated epitope was targeted during the first incubation, antibodies for the non-phosphorylated epitope was used in the second and vice versa. Importantly, as this technique did not involve removing the first primary antibody, antibodies from different hosts (mouse or rabbit) were used for phosphorylated and non-phosphorylated epitopes respectively. HRP inactivation did not affect the phosphospecific to non-phosphorylated signal ratios. For phospho-specific S6K1, we used two antibodies. The first antibody produced bands corresponding to ~80 kDa. This was slightly higher than expected (~70 kDa), though within the range defined by the manufacturer. Therefore a second antibody was used validate the results.

This antibody produced bands at a lower molecular weight (~60 kDa), corresponding to the predicted weight of the protein (UniProt identifier P23443-1). All incubation and washing steps were performed at 4°C using an automated membrane processor (Blot-Cycler, Precision Biosystems, Mansfield, MA, USA), except for S6K1-replication experiments, which was performed by hand in room temperature with incubations at 4°C. For each sample, total-protein and chemiluminescence quantification was calculated as the mean value of two separate experiments. Total-protein content was quantified using ImageJ (Rueden *et al*., 2017), and was defined as the mean gray value of the whole well with between-well values subtracted as background. Chemiluminescence signals were quantified using Image Studio Lite (LI-COR Biotechnology, Lincoln, Nebraska USA). Prior to statistical treatment, phospho-specific signals were normalized to the corresponding non-phosphorylated (pan-) signal from the same blot and pan-signals were normalized against the well total-protein content (Aldridge *et al*., 2008). In S6K1-replication experiment, phospho-specific signals were normalized to pan-signals using the total-protein stain to control for protein content between blots. Primary antibodies were purchased from Cell Signaling Technology (Leiden, The Netherlands): mTOR (Ser2448: #5536; pan: #4517), S6 kinase (Thr389 (∼80 kDa): #9206; Thr389 (∼60 kDa): #9234; pan: #2708), ribosomal protein S6 (Ser235/236: #4858; pan: #2317).

### Total RNA extraction, quantitative real-time reverse transcription polymerase chain reaction (qPCR) and mRNA sequencing

Approximately 25 *mg* of wet muscle-tissue was homogenized in a total volume of 1 ml of TRIzol reagent (Invitrogen, Life technologies AS, Oslo, Norway) using 0.5 mm RNase-free Zirconium Oxide beads and a bead homogenizer (Bullet Blender, Next Advanced, Averill Park, NY, USA) according to the manufacturer’s instructions. In order to enable analysis of target gene-expression per-unit tissue weight, an exogenous RNA control (*λ* polyA External Standard Kit, Takara Bio Inc, Shiga, Japan) was added at a fixed amount (0.04 *ng* × *ml*^−1^ of Trizol reagent) per extraction prior to homogenization, as previously described (Ellefsen *et al*., 2008, 2014*a*). Following phase-separation, 400 *µl* of the upper phase was transferred to a fresh tube and RNA was precipitated using isopropanol. The resulting RNA pellet was washed three times with 70% EtOH and finally eluted in TE buffer. RNA quantity and purity was evaluated using a spectrophotometer, all samples had a 260/280 *nm* ratio > 1.95. RNA was stored at −80°C until further processing. In the analysis of total RNA content per-unit tissue weight, one sample was excluded prior to analysis due to negative deviation from the expected value based on the relationship between sample weight and RNA content suggesting sample loss in washing steps. RNA integrity was assessed by capillary electrophoresis (Experion Automated Electrophoresis Station using RNA StdSens Assay, Bio-Rad) with average integrity scores (RQI) 8.1 (SD = 2.1).

Five-hundred nanograms of RNA were reverse transcribed using anchored Oligo-dT, random hexamer primers (Thermo Scientific) and SuperScript IV Reverse Transcriptase (Invitrogen) according to manufacturers instructions. All samples were reverse transcribed in duplicates and diluted 1:50 prior to real-time polymerase chain reaction (qPCR). qPCR reactions were run on a fast-cycling real-time detection system (Applied Biosystems 7500 fast Real-Time PCR Systems, Life technologies AS), with a total volume of 10 *µl*, containing 2 *µl* of cDNA, specific primers (0.5 *µM* final concentration) and a commercial master mix (2X SYBR Select Master Mix, Applied Biosystems, Life technologies AS). qPCR reactions consisted of 40 cycles (three seconds 95°C denaturing and 30 seconds 60°C annealing). Melt-curve analyses were performed for all reactions to verify single-product amplification. Gene-specific primers were designed for all targets using Primer-BLAST (Ye *et al*., 2012) and Primer3Plus (Untergasser *et al*., 2012) and ordered from Thermo Scientific, except for the external RNA control, for which primers were supplied with the kit. Raw fluorescence data was exported from the platform specific software and amplification curves were modelled with a best-fit sigmoidal model using the qpcR-package (Ritz & Spiess, 2008) written for R (R Core Team, 2018). Threshold cycles (Ct) were estimated from the models by the second-derivate maximum method with technical duplicates modeled independently. Amplification efficiencies were estimated for every reaction (as described by Tichopad *et al*., 2003; implemented in Ritz & Spiess, 2008). For every primer pair, mean amplification efficiencies (*E*) were utilized to transform data to the linear scale using *E*^−^*^Ct^*. Gene expression data was log-transformed prior to statistical analysis. As Ct-values, but not efficiencies are related to RNA integrity (Fleige & Pfaffl, 2006), RQI scores were used in the statistical treatment of qPCR data to control for potential degradation effects on a by target basis (see below).

### Data analysis and statistics

All descriptive data are presented as mean and standard deviation (mean (SD)) unless otherwise stated. To assess the effect of volume-conditions (number of sets) on muscle hypertrophy and strength, linear mixed-effects models were specified with relative changes from baseline as the dependent variable and number of sets as the main fixed effect. Baseline values were used as a co-variate together with sex. The interaction between sex and number of sets were explored for all hypertrophy and strength outcomes. Training-effects on molecular characteristics (Total-RNA and western-blot data) were also assessed using linear mixed-effects models specified with time and the time to exercise-volume interaction as fixed effects. Models were specified with random intercepts for participants and when appropriate, random slopes for time and exercise-volume on the level of participants. Model simplification was performed through reduction of random-effects parameters based on likelihood-ratio tests. Plots of residual and fitted values were visually inspected to assess uniformity of variance over the fitted range. Whenever deviations from these assumptions were identified, data were log-transformed and models were re-fitted.

Generalized linear mixed-effects models (GLMM) were used to fit muscle fibre distributions and gene-family normalized myosin heavy-chain mRNA data (Ellefsen *et al*., 2014*b*; after transformation to transcript counts as described by Matz *et al*., 2013) using the fixed and random effects structure specified above for molecular characteristics. A binomial variance/link-function (logit-link) was used for muscle fibre distributions with the number of counted fibres per sample used as weights to account for sample size. A beta variance/link-function (logit-link) was used to model gene-family normalized myosin heavy-chain mRNA data. This was done in order to account for the non-normal nature of relative fibre-type/myosin-isoform distribution data, where specific fibres/transcripts are analyzed as a proportion of the total number of fibers/transcripts in each sample and thus bound between 0 and 1. The beta model was used for gene-family mRNA data as the denominator could be regarded as arbitrary. Gene-abundance data, either expressed as per total-RNA or per-unit muscle weight using the external reference-gene were analyzed through modeling of gene-sets as suggested by Matz *et al*. (2013) using mixed linear models with within-model normalization through the addition of random effects of technical replicates. To allow for gene-specific variances, variance functions were specified per strata (per gene) (Pinheiro & Bates, 2000). RNA integrity scores (RQI) were included in the model on a per target basis to control for RNA degradation.

Tests against the null-hypotheses of no differences between volume-conditions and no effect of time were performed on model-parameter estimates resulting from LMM and GLMM. LMM were fitted using the nlme-package (Pinheiro & Bates, 2000), binomial GLMM models using the lme4-package (Bates *et al*., 2015) and beta GLMM using glmmTMB-package (Magnusson *et al*., 2019) written for R.

To explore determinants of additional benefit of multiple-sets, dichotomous response variables were constructed from individual differences in single- and multiple-sets outcomes in muscle-hypertrophy (CSA), knee-extension and leg-press 1RM. When the difference between volume-conditions in training-induced outcomes were larger than the estimated measurement error in the direction of multiple-sets, variables were coded as additional benefit of multiple-sets. The measurement error was estimated from the baseline between-legs coefficient of variation (CV). The probability of additional benefit of multiple-sets was related to a wide range of predictors using logistic regression. Prior to model fitting, a-priori selection of relevant predictor variables were done, these included blood variables, baseline strength and muscle mass, volume-dependent molecular responses to training (i.e. total-RNA content and mTOR pathway phosphorylation expressed as a percentage of single-sets readouts) and baseline fibre-type composition. Purposeful selection of variables were done in a step-wise manner following (Hosmer *et al*., 2013), first each possible predictor was fitted in univariate models and predictors with *P* < 0.20 were kept for further considerations. All predictors from the first step was fitted in a preliminary model from where predictors were sequentially removed if they were not significant at the *P* < 0.1-level using Wald-based *P*-values or influenced other predictors 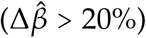. As a last step, predictors removed in the first step was fitted to the reduced model and the model was reduced to the final formulation. Logistic models fitted with small samples has been shown to give biased estimates (Nemes *et al*., 2009), this was recognized and bias-corrected estimates were reported (Kosmidis, 2019) with *P*-values from likelihood-ratio tests comparing sequentially reduced models.

The level of statistical significance was set to *α* = 0.05. All data-analysis was done in R (R Core Team, 2018).

## Results

### Volume-dependent regulation of muscle strength, muscle mass and fiber type composition

Overall, 12 weeks of resistance training led to 46% (95% CI: [39, 53], *P*<0.001) increase in muscle strength (1RM) and 4.4% ([3.2, 5.6], *P*<0.001) and increase in muscle mass when averaged over volume-conditions. Adherence to the protocol was 96 (5)% of the precribed 31 sessions (range 81-100%), which gives an efficiency for developing muscle strength and mass equivalent to 1.60 (0.64)% and 0.15 (0.12)% per session, being within the expected range of training-induced changes (Ahtiainen *et al*., 2016).

Training had no effect on serum levels of cortisol and testosterone (Table 2). IGF-1 decreased ~5.4 % from Week 0 to Week 2, and increased ~3.6 % from pre- to post-exercise in Week 2. Growth hormone concentrations increased in response to acute exercise, with patterns differing between sexes (Table 2). Vitamin D levels were different at baseline between males (76.6 (16.4) *nmol* × *L*^−1^) and females (100.0 (33.4) *nmol* × *L*^−1^, *P* = 0.006) and were similarly reduced from Week 0 to Week 12 in both sexes (63.1 (19.8) and 91.4 (31.7) *nmol* × *L*^−1^ for males and females respectively, time-effect *P* < 0.001).

**Table 2:**
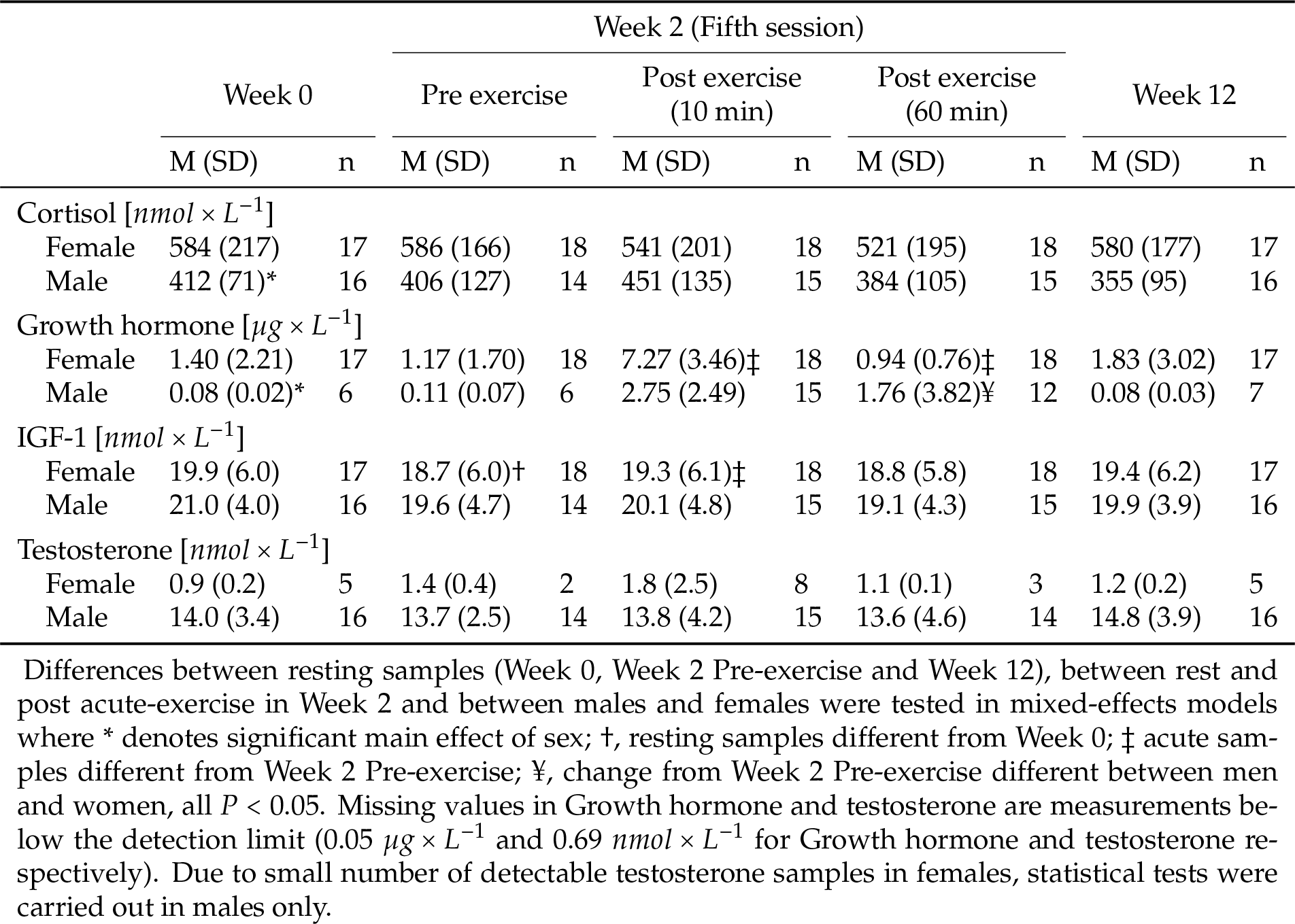
Hormone measurements

The difference in number of sets per exercise between multiple- and single-set conditions resulted in a ratio of performed work (number of repetitions × external resistance) between legs corresponding to 2.9 (0.3) in knee extension and 3.0 (0.5) in leg press. This was accompanied by higher ratings of perceived exertion in response to multiple sets than single sets (7.09 (1.95) vs. 6.22 (1.82), *P* < 0.001). Concomitantly, multiple-set resistance-training led to greater increases in muscle strength over the course of the intervention than single-set training (all variables *P* < 0.05, Figure 2C). This difference appeared late in the intervention for both leg press (1RM, after nine weeks) and leg extension (1RM, after twelve weeks, Figure 2D). In line with this, multiple-sets training led to greater increases in knee extensor CSA (mean percentage-point difference 1.62, [0.75, 2.50], *P* <0.001, Figure 2B). There was no difference between sexes in relative muscle strength and mass gains, and sex did not interact with responses to different volume conditions. There was a strong correlation between responses to multiple-sets and single-set conditions with respect to both 1RM strength gains (knee-extension, *r* = 0.88, [0.77, 0.94], *P* < 0.001; leg-press, *r* = 0.91, [0.82, 0.96], *P* < 0.001, Figure 6A) and muscle mass (*r* = 0.75, [0.55, 0.87], *P* < 0.001, Figure 6B). Increases in muscle 1RM strength correlated with increases in mass (*r* = 0.39, [0.06, 0.64], *P* = 0.023, Figure 2E) assessed as averaged effects of the two volume conditions.

In muscle tissue, multiple-sets training led to more pronounced conversion of Type IIX fibres into Type IIA fibres from Week 0 to Week 12 than single-set training, measured as both cell counts using immunohistochemistry (OR: 0.53, [0.30, 0.92], Figure 5B) and mRNA abundance using gene-family profiling (OR: 0.76, [0.63, 0.92], Figure 5B). Surprisingly, at week 2, the relationship between training volume and fiber conversion was the opposite, with single-set legs showing greater IIX to IIA transition (OR: 1.60, [1.04, 2.48]. Notably, from baseline to week 2, a pronounced decrease was seen in MYH1 gene expression (coding for the Type IIX myosin-heavy chain transcript) and more so in response to multiple-sets training than to single-set training. This change that was partly reversed in week 12 (Figure 5C).

### Volume-dependent regulation of mTOR-signaling and ribosomal biogenesis

Multiple-sets training led to greater phosphorylation of mTOR, S6K1 and rpS6 than single-sets training (Figure 3A), measured in muscle biopsies sampled after the fifth training session (mean %-difference from single-sets with [95% CI]: phospho-mTOR, 11.8 [2.5, 22.1], phospho-S6K1, 19.1 [0.3, 41.4]; phospho-rpS6, 28.4 [4.7, 57.4]). For S6K1, this was confirmed using a separate antibody aimed at the same phosphorylation-site but producing quantifiable bands at a slightly lower molecular weight (~60 vs. ~80 kDa) (58.8 [13.7, 121.9]%, Figure 3C-E). Together this suggests volume-dependent regulation of the mTOR-pathway. Compared to baseline, non-phosphorylated (pan-) levels of mTOR (pan-mTOR) increased at all time-points (Week 2 Pre-ex, 9.4 [3.9, 15.1]; Week 2 Post-ex, 11.5 [5.5, 17.8]; Week 12, 6.0 [0.2, 12.1]), pan-levels of rpS6 increased at all rested-state biopsy time-points (Week 2 Pre-ex, 22.0 [8.0, 37.9]; Week 2 Post-ex, −18.3 [−29.6, −5.2]; Week 12, 14.7 [1.4, 29.8]), and pan-levels of S6K1 remained unchanged at all rested-state biopsy time-points. There were no effects of training volume on non-phosphorylated protein abundances.

**Figure 3:**
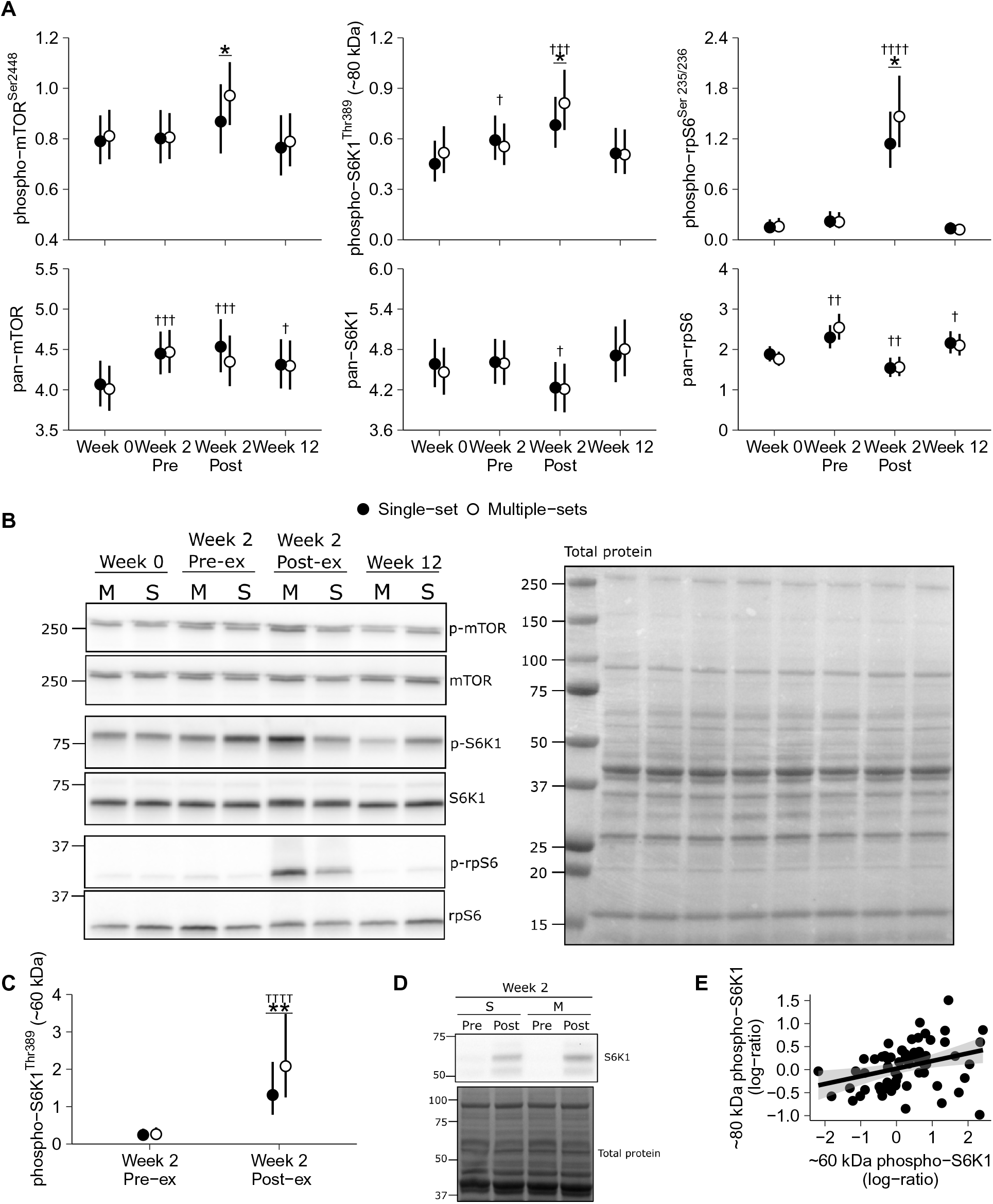
Western-blot analysis of the mTOR-signaling pathway. Training-volume dependent phosphorylation of mTOR, S6K1 and rp-S6 proteins in m. Vastus lateralis measured after single bouts of multiple-(M) and single-set (S) resistance training at Week 2 (A). Representative blots and total-protein stains are shown in B and D. Phospho-S6K1 were measured using two antibodies (A, original analysis; C-D, supplementary analysis; see Methods), with multiple- vs single-set signal ratios correlating between the two (E, Spearman’s *ρ* 0.40, *P* < 0.001). Values are mean values ± 95% CI. Points represents log-ratios of volume-conditions (E). † represents difference from Week 0 †-†††† for *P* < 0.05 - *P* < 0.0001; * represents differences between volume conditions, * - * * for *P* < 0.05 - *P* < 0.01.

In line with these data, multiple-sets training resulted in 8.8 [1.5, 16.6]% greater total RNA abundance per-weight-unit muscle tissue at Week 2 than single-set training. This difference was also evident at Week 12, albeit less extensive (5.9 [−1.0, 13.3]%, Figure 4A). Accordingly, the multiple-sets leg showed greater abundances of mature rRNA transcripts at Week 2, particularly rRNA 18S (18S, 19.4 [0.8, 41.4]%; 28S, 14.5 [−1.8, 33.6]%; 5.8S 14.7 [−1.20, 33.18]%). The abundances of these rRNA subspecies remained elevated at week 12, though without clear differences between volume conditions (Figure 4B). The rRNA precursor transcript 45S, measured per-unit total-RNA, did not increase from baseline to week 2, but increased by 48.8 [3.6, 113.6]% in the single-sets condition at week 12, with multiple sets remaining near baseline levels (−28.8 [−50.4, 2.1]% of single sets). Overall, these data suggest that resistance training-induced increases in ribosomal content depend on training volume. Further supporting this view, mRNA expression of the transcription factor c-Myc, which is important for initiating rRNA transcription (Riggelen *et al*., 2010), increased 1.58 [1.14-2.17]-fold more in response to multiple-sets training than to single-set training (Figure 4C, measured before and after the fifth training session).

**Figure 4:**
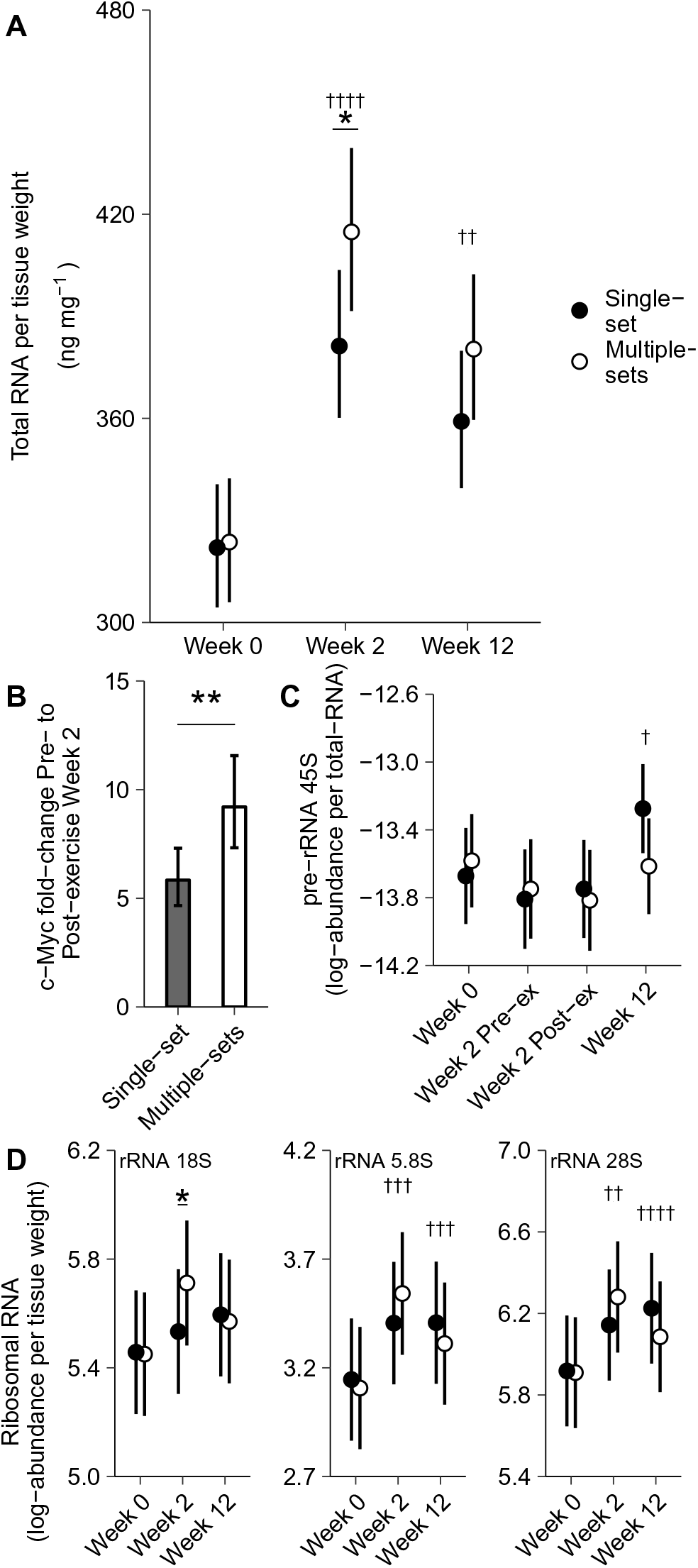
Total-RNA and ribosomal RNA. Training-volume dependent changes in total RNA in m. Vastus lateralis after 2 weeks of resistance training (measured per-unit muscle weight, Week 2, A), c-Myc mRNA measured 1h after a training session at Week 2 (B) and ribosomal RNA 18S at Week 2 (D). Other mature ribosomal RNA species exhibited similar expression patterns without reaching statistical significance (D). Ribosomal pre-RNA 45S expressed relative to total RNA showed greater relative abundances at Week 12 than Week 0 in the single-set leg (C). Values are estimated means ± 95% CI. * represents difference between volume conditions for *P* < 0.05. † represents difference from Week 0, †-†††† for *P* < 0.05 - *P* < 0.0001.

### Determinants of additional benefit of multiple-sets training

Fifteen participants showed a robust benefit of multiple-sets over single-sets for increases in CSA, determined as differences in training-induced changes greater than the average baseline between-leg variation in favour of multiple-sets (2.4% between leg variation at baseline, Figure 5A). To identify determinants of multiple-set benefits, we performed logistic regression analyses with purposeful selection of variables. Variables initially selected for modelling are listed in Table 3. After variable selection, total RNA content per-unit tissue weight measured at rest in Week 2 remained as the single predictor (Table 4), with total RNA content being greater in the group having robust benefits of multiple sets (Figure 5C). For every percentage-point increase in total-RNA in the multiple-sets leg (compared to the single-set leg), the odds of multiple-sets benefit increased by 1.05 [1.00, 1.11] (Table 4). In all models, sex was included as a calibrating variable to account for potential predictors with sex-dependent regulation (e.g. blood variables). However, excluding sex and apparent sex-dependent variables from the variable selection, did not affect the conclusion. As for muscle strength, 18 and 15 participants showed benefits of multiple sets for increases in 1RM knee-extension and leg-press (defined as a difference in training-induced changes in favour of multiple-sets greater than the baseline between-leg variation, 2.9 and 4.0% for the knee-extension and leg-press 1RM respectively). Variable selection-analyses did not reveal significant determinants for this phenomenon.

**Figure 5:**
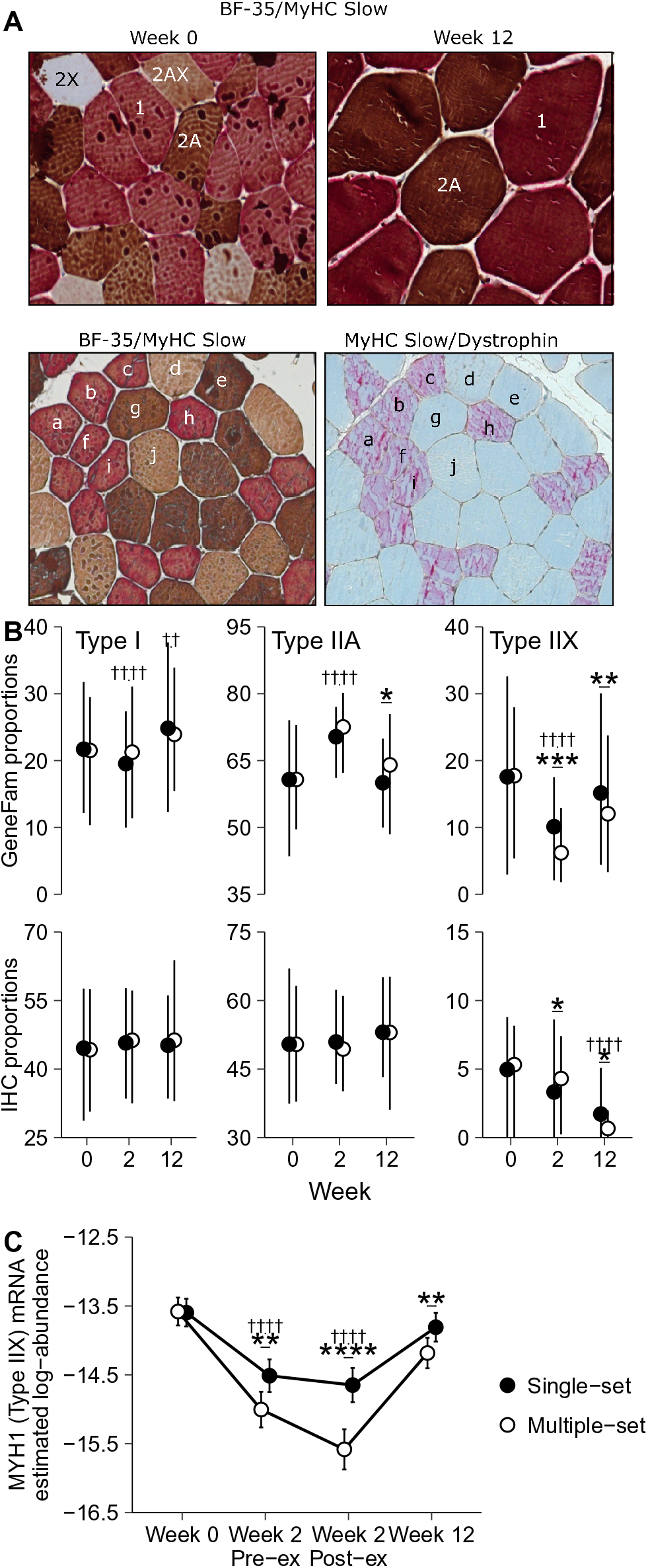
Fiber-type distributions. Volume-dependent changes in muscle fibre-type distribution in m. Vastus lateralis after 2 and 12 weeks of multiple and single-set resistance training, measured as relative cell counts using immunohistochemistry (A and B) and gene family profiling (GeneFam)-normalized myosin heavy-chain mRNA expression (C). The volume-dependency was evident as surplus reductions in Type IIX mRNA abundance at all time points (MYH1, D). Values are mean ± 10 − 90*^th^* percentile in B and mean ± 95% CI in C. † represent difference from Week 0, †-†††† for *P* < 0.05 - *P* < 0.0001; * represent differences between sets * - * * * * for *P* < 0.05 - *P* < 0.0001.

**Figure 6:**
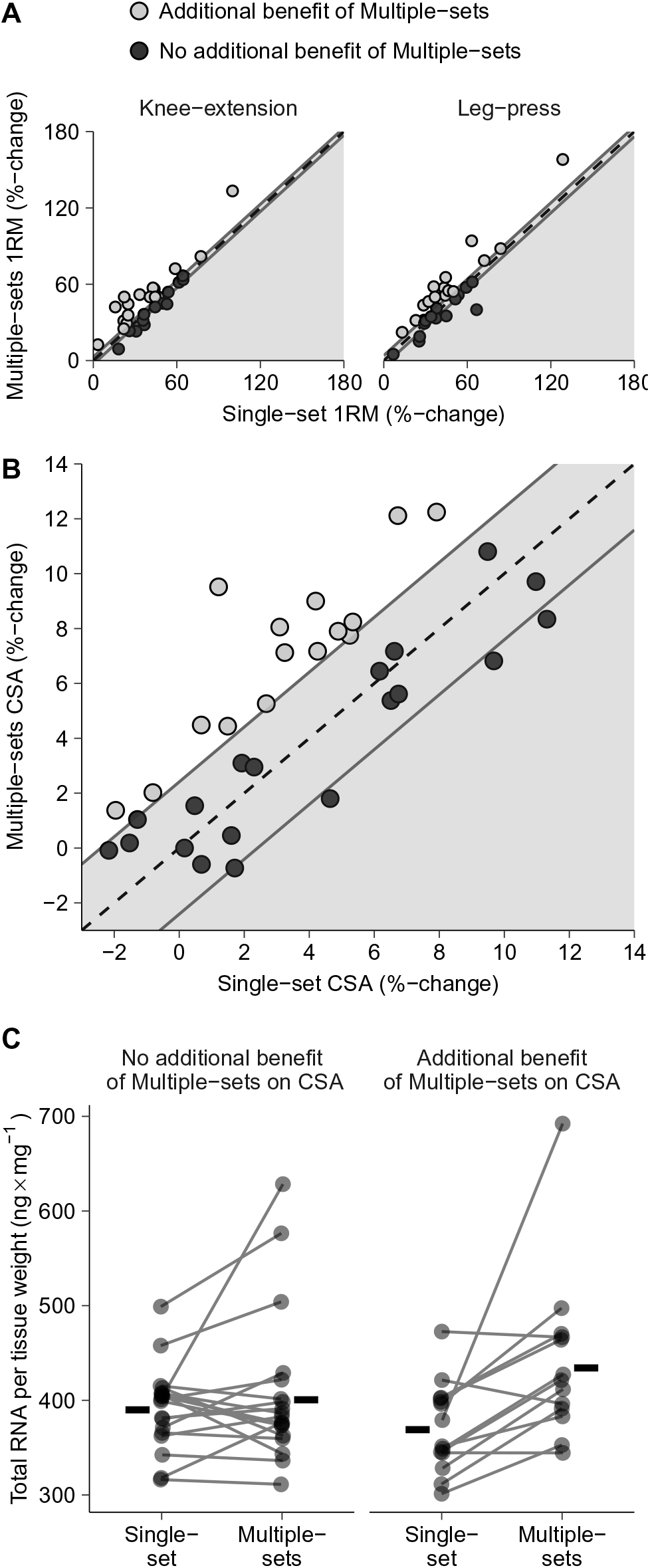
Strength (A) and Hypertrophy (B) responses and total RNA grouped according to benefits of multiple sets. Participants that showed additional benefit of multiple-sets on training-induced muscle hypertrophy (B) displayed higher total RNA content in m. Vastus lateralis after two weeks of training (C) (interaction Benefit × Sets *P* = 0.015). Strength and hypertrophy responses to multiple- and single-set training showed large correlation (knee-extension, *r* = 0.88 95% CI: [0.77, 0.94], *P*<0.001; leg-press, *r* = 0.91 [0.82, 0.96], *P*<0.001, A; and muscle mass, *r* = 0.75 [0.55, 0.87], *P*<0.001, B. Dashed lines in A and B is the identity line (*y* = *x*), the distance from dashed to solid lines represent the baseline between-leg variation. Horizontal lines in C represents group means, connected points represents individual values.

**Table 3:** Logistic regression coefficients for additional benefit of Multiple-sets on training-induced hypertrophy

| Variable | Mean (SD)<sup>a</sup> |  | Logistic regression-coefficients<sup>b</sup> |  |  |  |
| --- | --- | --- | --- | --- | --- | --- |
|  | ♀ | ♂ | Odds-ratio | 95% CI | Deviance | P-value<sup>c</sup> |
| **Ribosome biogenesis** |  |  |  |  |  |  |
| Total-RNA Week 2 (% of single-sets) | 15 (22) | 3.3 (14) | 1.05 | [1.00, 1.11] | 6.70 | 0.010 |
| Total-RNA Week 12 (% of single-sets) | 3.8 (18) | 12 (20) | 1.01 | [0.97, 1.04] | 0.11 | 0.735 |
| **mTOR signaling** |  |  |  |  |  |  |
| mTOR<sup>Ser2448</sup> (% of single-sets) | 14 (25) | 25 (68) | 1.00 | [0.98, 1.01] | 0.21 | 0.647 |
| S6K1<sup>Thr389</sup> (% of single-sets) | 42 (62) | 29 (79) | 1.00 | [0.99, 1.01] | 0.17 | 0.678 |
| rpS6<sup>Ser235/236</sup> (% of single-sets) | 78 (123) | 26 (47) | 1.00 | [0.99, 1.01] | 0.02 | 0.879 |
| **Blood parameters** |  |  |  |  |  |  |
| Vitamin D (Week 0) | 100 (33) | 77 (16) | 0.99 | [0.96, 1.01] | 1.38 | 0.241 |
| Testosterone (Mean Week 0-2)<sup>d</sup> | 0.70 (1.0) | 14 (2.9) | 0.73 | [0.48, 1.11] | 3.81 | 0.051 |
| IGF-1 (Mean Week 0-2) | 19 (5.5) | 20 (4.2) | 1.04 | [0.90, 1.20] | 0.38 | 0.540 |
| Cortisol (Mean Week 0-2) | 570 (164) | 419 (71) | 1.00 | [1.00, 1.01] | 0.28 | 0.595 |
| Growth hormone (Week 2 Post-ex) | 7.3 (3.5) | 2.7 (2.5) | 1.02 | [0.81, 1.29] | 0.04 | 0.838 |
| **Muscle fibre-types<sup>e</sup>** |  |  |  |  |  |  |
| Type 2A (% of total MHC) | 49 (6.0) | 51 (9.4) | 0.99 | [0.91, 1.08] | 0.05 | 0.827 |
| Type 2X (% of total MHC) | 5.0 (6.1) | 4.0 (2.4) | 1.18 | [0.97, 1.44] | 4.98 | 0.026 |
| Type 1 (% of total MHC) | 46 (9.4) | 45 (9.4) | 0.97 | [0.90, 1.05] | 0.82 | 0.365 |
| **Baseline characteristics** |  |  |  |  |  |  |
| Baseline Leg extension 1RM (kg<sup>-1</sup>) | 0.78 (0.15) | 0.99 (0.09) | 1.99 | [0.031, 126] | 0.13 | 0.721 |
| Baseline Leg press 1RM (kg<sup>-1</sup>) | 2.4 (0.58) | 2.8 (0.76) | 0.62 | [0.273, 1.40] | 1.73 | 0.188 |
| Baseline lean mass (%) | 65 (5.9) | 80 (5.8) | 1.09 | [0.96, 1.23] | 2.11 | 0.147 |
| **Training characteristics** |  |  |  |  |  |  |
| Total number of sessions | 30 (1.7) | 30 (1.5) | 0.96 | [0.63, 1.46] | 0.05 | 0.824 |
| Supervised sessions | 92 (8.3) | 90 (11) | 0.96 | [0.89, 1.04] | 1.22 | 0.269 |

<sup>a</sup> Descriptive statistics are grouped by sex; <sup>b</sup>, Sex was kept as a covariate in all models to account for sex-differences in independent variables; <sup>c</sup>, P-values are derived from likelihood-ratio tests; <sup>d</sup> testosterone measurements below detection limit coded as 0; <sup>e</sup>, baseline average of both legs.

**Table 4:** Multiple logistic regression models on additional benefit of multiple-sets on training-induced hypertrophy.

| Variable | Estimate<sup>a</sup> | SE | Z-value | Wald *P*-value | LRT<sup>b</sup> *P*-value |
| --- | --- | --- | --- | --- | --- |
| **Model 1** |  |  |  |  |  |
| Intercept | -8.43 | 5.53 | -1.53 | 0.127 |  |
| Sex (Male) | 1.26 | 3.07 | 0.41 | 0.682 |  |
| Total-RNA Week 2 (% of single-sets) | 0.05 | 0.03 | 1.47 | 0.140 |  |
| Testosterone (Mean Week 0-2) | -0.14 | 0.21 | -0.67 | 0.502 |  |
| Type 2X (% of total MHC) | 0.12 | 0.10 | 1.23 | 0.219 |  |
| Baseline lean mass (%) | 0.10 | 0.08 | 1.27 | 0.202 |  |
| **Model 2** |  |  |  |  |  |
| Intercept | -8.48 | 5.39 | -1.57 | 0.116 |  |
| Sex (Male) | -0.56 | 1.40 | -0.40 | 0.688 |  |
| Total-RNA Week 2 (% of single-sets) | 0.05 | 0.03 | 1.56 | 0.118 | Model 1 vs 2 *P* = 0.381 |
| Type 2X (% of total MHC) | 0.13 | 0.10 | 1.33 | 0.184 |  |
| Baseline lean mass (%) | 0.10 | 0.08 | 1.28 | 0.202 |  |
| **Model 3** |  |  |  |  |  |
| Intercept | -1.84 | 0.81 | -2.28 | 0.023 |  |
| Sex (Male) | 0.89 | 0.85 | 1.05 | 0.294 |  |
| Total-RNA Week 2 (% of single-sets) | 0.05 | 0.03 | 1.73 | 0.083 | Model 2 vs. 3 *P* = 0.144 |
| Type 2X (% of total MHC) | 0.13 | 0.10 | 1.30 | 0.192 |  |
| **Model 4** |  |  |  |  |  |
| Intercept | -1.23 | 0.64 | -1.91 | 0.056 |  |
| Sex (Male) | 0.76 | 0.82 | 0.92 | 0.357 | Model 3 vs. 4 *P* = 0.078 |
| Total-RNA Week 2 (% of single-sets) | 0.05 | 0.03 | 1.91 | 0.057 |  |
| **Model 5** |  |  |  |  |  |
| Intercept | -0.43 | 0.48 | -0.89 | 0.375 | Model 4 vs. 5 *P* = 0.010 |
| Sex (Male) | 0.16 | 0.72 | 0.22 | 0.826 |  |

<sup>a</sup>, Estimates are log-odds ratios; <sup>b</sup>, P-values derived from Likelihood ratio test were used for inference

## Discussion

In the present study, multiple-set resistance training led to greater increases in muscle strength and mass than single-set training. This is in agreement with results from meta-analyses concluding in favor of moderate-compared to low-volume training (Krieger, 2009, 2010; Schoenfeld *et al*., 2016). The greater effect of multiple-sets training coincided with greater responses in muscle biological traits indicative of hypertrophic response (Andersen & Aagaard, 2000; Goodman *et al*., 2011; Terzis *et al*., 2008; Luo *et al*., 2019; Stec *et al*., 2016), including greater transition from Type IIX to IIA muscle fibres, greater post-exercise phosphorylation of mTOR, S6-kinase and ribosomal protein S6, greater post-exercise expression of c-Myc and greater rested-state levels of total RNA and ribosomal RNA. While most of these variables are already assumed to be volume sensitive, such as muscle mass and strength (Krieger, 2009, 2010; Schoenfeld *et al*., 2016) and mTOR-signaling (Burd *et al*., 2010; Terzis *et al*., 2010), this is the first study to suggest that the IIX → IIA fiber switch is also volume sensituive. Importantly, this adaptation is a hallmark of resistance training adaptations (Andersen & Aagaard, 2000). This study also suggests that the volume-sensitive increase in ribosomal content is essential for beneficial effects of increases in training volume on muscle growth, as shown by fifteen of the participants. Arguably, the biological resolution of the present data was high due to the use of a within-participant training model, facilitating disclosure of volume-dependent effects. Indeed, previous studies have typically used between-participants models to assess the volume-dependency of muscle development (e.g. Starkey *et al*., 1996; Ronnestad *et al*., 2007; Rhea *et al*., 2002) or have failed to account for the within-participant perspective in their analyses (Mitchell *et al*., 2012). This makes their interpretations prone to the large individual-to-individual variation in exercise adaptability (seen in e.g. Ahtiainen *et al*., 2016), which has been linked to variation in genetic and epigenetic predisposition (Timmons, 2011; Seaborne *et al*., 2018), and may potentially explain the long-standing lack of consensus (Carpinelli & Otto, 1998; Krieger, 2010).

In the present study, a large span of inter-individual variation in training responses was evident for both gains in muscle strength and muscle mass. The observed variation in muscle hypertrophy (SD of average %Δ CSA ~4%) was comparable to that seen in larger cohorts (Ahtiainen *et al*., 2016). The strong correlation between responses to the two volume-conditions (see Figure 6A) further highlights the importance of within-participant analyses: if responses to one training protocol were strong, responses to the other protocol were also strong. Consequently, our contralateral protocol resulted in lower estimates of differences between volume-conditions on the population level, expressed as relative gains in muscle mass per week, compared to a previous meta-analysis (~1.6 vs. ~2.5% estimated from Table 3 in Schoenfeld *et al*., 2016). Notably, in the present study, this comparison was prone to systemic contralateral adaptions to training, which would diminish differences between volume conditions. However, this effect is likely negligible as non-trained limbs typically do not show increased protein synthesis, hypertrophy or muscle fibre type transitions (Brook *et al*., 2016; Wilkinson *et al*., 2006). Instead, it is plausible that the overall effect of added training-volume reported in (Schoenfeld *et al*., 2016) is overestimated due to small sample sizes, a known weakness in meta analyses (Nüesch *et al*., 2010). Thus, contralateral designs arguably provide more accurate comparisons of responses to different training volumes on the population level, accounting for inter-individual differences in responses.

In our search for determinants that could explain the variation in muscle growth patterns to the two volume protocols, potential explanatory factors included baseline characteristics, blood variables, indices of mTOR-signaling and ribosome biogenesis as well as training charactersitics. Following variable selection, the only variable that could explain additional benefits of multiple-over single-set training was levels of total RNA at week 2 in the multiple-sets leg. As total RNA is a valid proxy marker of rRNA abundance (Zak *et al*., 1967; Chaillou *et al*., 2014), this suggests that early-phase, volume-dependent ribosomal accumulation is a determinant of dose-response relationships between training volume and muscle hypertrophy. In other words, the ability to induce superior increases in ribosomal content in response to higher training volume is necessary to induce subsequent superiority in growth, probably acting by increasing protein synthesis capacity. This fits well with the overall impression conveyed by the data set, wherein multiple-sets training resulted in larger increases in total RNA and mature rRNA species (rRNA 18S, 28S and 5.8S). In untrained participants, early accumulation of ribosomal content seems to be a generic response to training (Brook *et al*., 2016; Stec *et al*., 2016). This accumulation follows a progressive nature during the first three weeks of training (Brook *et al*., 2016) whereupon total RNA remains at elevated levels for at least 12 weeks (Figueiredo *et al*., 2015; Mobley *et al*., 2018, 2018), assumingly preceded by increased expression of the 45S pre-rRNA. The latter was not evident in the present data, suggesting that timing of muscle biopsy-sampling was not suited for investigating *de novo* synthesis of rRNA. The potential link between ribosomal content in muscle and trainability is not surprising. Several studies have shown that ribosomal biogenesis measured as total RNA per tissue weight is positively associated with training induced muscle hypertrophy (Stec *et al*., 2016; Figueiredo *et al*., 2015; Mobley *et al*., 2018) in addition to early observations of a relationship between RNA content and rate of protein synthesis(Millward *et al*., 1973).

Variable selection did not identify other variables that could explain benefits of moderate training volume, discarding biological traits such as sex, baseline values of lean mass and muscle fiber composition. Variable selection also discarded phosphorylation of mTOR, along with phosphorylation of its downstream targets. This seems somewhat counterintuitive, as these signaling cues are regulators of ribosomal biogenesis and function (Nader *et al*., 2005; Riggelen *et al*., 2010; West *et al*., 2016), giving them potential roles in accumulation of rRNA and total RNA and moderate-volume beneficence. However, these signaling cues are acute-phase responders to resistance training that show phasic and time-dependent regulation. This means that the measured change in for example mTOR phosphorylation depends on factors such as timing of biopsy sampling, giving it low resolution power and making it less suited for explanatory analyses. In accordance with this, the association between acute mTOR signaling and hypertrophy in humans is ambiguous in the literature, with some studies showing correlations with degrees of muscle hypertrophy (Terzis *et al*., 2008; Mitchell *et al*., 2013) while others do not (Mitchell *et al*., 2012; Phillips *et al*., 2017). Obviously, this does not mean that the volume-dependent phosphorylation of mTOR and its targets was without a role in the observed RNA response patterns. It simply means that we were not able to detect any such association. Whereas training-induced mTORC1 activity is transitory, its effects are long lasting, leading to chronic adaptations such as accumulation of ribosomal RNA, which are easily detected in rested muscle. Targeting such rested-state muscle characteristics obviates issues such as biopsy-sampling timing, making them better suited as biomarkers. In addition, the role of mTORC1 in ribosomal biogenesis is likely synergistic and includes parallel pathways such as induction of c-Myc and its downstream targets (West *et al*., 2016)

Initially, we hypothesized that participants with lower proportions of Type IIX muscle fibers would benefit more from moderate volume training (and vice versa) than subjects with higher proportions of IIX, as outlined in the pre-study clinical trials registration. This hypothesis was rooted in prevailing training guidelines, advocating higher training volume for individuals with lesser training experience (and thus likely lower proportions of IIX fibres) (Ratamess *et al*., 2009). Indeed, during variable selection, baseline IIX fibre proportions were selected as one potential explanatory factors behind volume benefits on hypertrophy (Table 3). However, contrary to our hypothesis, higher levels of IIX tended to explain beneficial effects of multiple sets. Although this trait was discarded from the final model, the tendency towards a positive effect of higher IIX levels could be ascribed to their greater growth potential (Stec *et al*., 2016; Jespersen *et al*., 2011), with these fibres having been in a state of disuse prior to the intervention. This implies a relatively rapid transition of type IIX fibres into IIA fibres, which indeed was present in the data already after two weeks of training at both protein and RNA levels. Correlation analyses revealed that this transition was more pronounced in individuals with higher baseline levels of IIX, with an *r*-value >0.95 (data not shown), far exceeding the bias expected from regression-towards-the-mean.

To our knowledge, this is the first study to show that muscle fibre transitions from Type IIX to IIA depend on resistance training volume. Moderate volume resulted in 1.5%-point greater reductions in Type IIX fibre expression from baseline to post intervention compared low volume, presumably driven by more pronounced reductions in mRNA expression of the MYH1 (Myosin heavy chain IIX) gene (−61% vs. −31%). Previous studies have not compared this transition directly between volume protocols. However, Pareja-Blanco *et al*. (2017) observed blunted IIX → IIA transitions in response to non-exhaustive high-load resistance training compared to load-matched training to volatile failure. Together with our data, this makes exercise volume and subsequent metabolic stress and dosage of neuromuscular activity plausible candidates for regulation of IIX → IIA reprogramming, as opposed to mechanical stimuli. Indeed, in rodents, mechanical load does not affect fibre-type transitions (Eftestol *et al*., 2016), which is instead linked to neural activation. Interestingly, after 2 weeks of training, the volume effect on IIX → IIA transitions was opposite to our main finding after 12 weeks, with low-volume resistance training resulting in more pronounced decreases on the cell level. This was not evident at the mRNA level, as moderate volume showed distinct benefits also at this time point, with heavily suppressed levels of MYH1 mRNA. Whether these discrepancies are due to increased need for tissue-repair in the moderate-volume leg at two weeks (Kim *et al*., 2005; Damas *et al*., 2016) or other causalities, rather than myofibril-specific adaptations remain unclear. Regardless of causality, these data underline the importance of optimizing exercise volume to achieve optimal training progression, such as by making use of progressive volume protocols. Such protocols remain largely unexplored, but it seems evident that in the untrained, too large or too small training volumes in the first phase of a training intervention may lead to suboptimal adaptations.

In conclusion, resistance training with higher volume led to surplus increases in muscle CSA, muscle strength and fibre-type transitions, as well as greater responses in molecular hypertrophy signaling and effectors. Beneficial effects of multiple-sets over single-set training on muscle hypertrophy coincided with higher total RNA levels at week 2, suggesting that volume-dependent early-phase regulation of ribosomal biogenesis determines the dose-response relationship between training volume and muscle hypertrophy.

## Additional information

### Competing interests

No conflicting interests.

### Author contributions

Data collection was done in the Sport Science Laboratory at Inland University of Applied Sciences and the Hospital for Rheumatic Diseases with molecular analyses partly performed at Åstrandlaboratoriet, The Swedish School of Sport and Health Sciences and Innlandet Hospital Trust. DH, SE, BRR designed the study; DH, SJØ, LK, MH, SE and WA performed experiments; DH analysed the data; DH and SE interpreted the results; DH drafted the manuscript; DH, SJØ, LK, MH, BRR, EB, WA, JEW, IH and SE edited and revised the manuscript. All authors have approved the final version of the manuscript and agree to be accountable for all aspects of the work. All persons designated as authors qualify for authorship, and all those who qualify for authorship are listed.

### Funding

The work presented here was supported by a grant from Innlandet Hospital Trust (grant nr. 150282).

## Acknowledgements

The authors would like to express their gratitude to Johanne Seeberg, Stine Dahl, Marianne Bratlien, Martin Nordseth, Erlend Hakestad, Ole-Martin Hveem and Mari Skifjeld for their skilled assistance and commitment in performing the training intervention. The authors are also grateful for the technical support of Håvard Nygaard, Gunnar Slettaløkken Falch, Jostein Flata, Bente Malerbakken and Anne Grete Mathisen. The study was made possible through the effort of all dedicated participants, the authors are humbly impressed at thankfull for their contribution.

